## Supporting Information for "The macroecological dynamics of sojourn trajectories in the human gut microbiome"

**1 Quantitative Life Sciences, The Abdus Salam International Centre for Theoretical Physics (ICTP), Trieste, Italy.**

*** Contact:** ``

### Data

Below we describe the processing pipeline for temporal sequence data from human hosts as well as how the three empirical sojourn trajectory patterns were identified. We note that, to our knowledge, this is the first instance of sojourn trajectory patterns being investigated in any ecological system.

#### Data processing

Human gut microbiome timeseries from 16S rRNA amplicon datasets were reprocessed to ensure standardization. We focused on datasets where hosts were sampled for a minimum of 100 days ([1, 2, 3]; Table S1). FASTQ files were downloaded and reprocessing was performed using `DADA2 v1.16` [4]. Each dataset was reprocessed using the flag `pool=TRUE` so that ASVs with a read count of one in a given community (i.e., singletons) could be inferred, allowing us to examine the entirety of the empirical sampling distribution. Taxonomy was assigned using `v138.1` of the Silva non-redundant (NR) training set at 99% similarity. Species-level taxonomy was assigned using `v138.1` of the Silva species-level assignment dataset.

We elected to split the timeseries of two hosts from one dataset to account for perturbations to the microbiome [3]. Specifically, for host A we split the timeseries into pre (0 - 70 days) and post-travel (123 - 364 days) periods. Host B acquired a *Salmonella* infection, requiring us to split the timeseries between pre (0 - 150 days) and post-infection (160 to 252 days) periods. The macroecological consequences of this disruption were previously investigated [5].

We investigated the time between sampling events ($\delta t$) for each host in each dataset. We found that several hosts in Poyet et al. had a low number of consecutive samples with high $\delta t$ ($\sim$ 60 days). Such extremely long times between sample collections can make it difficult to investigate the temporal dynamics as they greatly exceed the expected timescale of growth, having the potential to confound one's interpretation of comparisons between empirical and time-permuted null distributions [6]. Therefore, we elected to split a timeseries into two if there was a sampling interval $\delta t > 20$. The resulting split timeseries had roughly even sampling intervals and we kept those with $\geq 60$ observations, resulting in the removal of hosts ae and an from Poyet et al. [2].

#### Data analysis

We start with the observation that the distributions of relative abundances of a given ASV over time tends to follow a gamma distribution, known as the Abundance

Fluctuation Distribution (AFD).

$$P\left(x_i|\beta_i, \frac{\beta_i}{\bar{x}_i}\right) = \frac{1}{\Gamma\left(\beta_i\right)} \left(\frac{\beta_i}{\bar{x}_i}\right)^{\beta_i} \exp\left[-x_i \frac{\beta_i}{\bar{x}_i}\right] x_i^{\beta_i - 1} \tag{S1}$$

where $\bar{x}_i$ is the mean relative abundance and $\beta_i$ is the inverse squared coefficient of variation. In order to account for the number of reads as a form of sampling, we assume that sampling occurs as the Poisson-limit of a multinomial sampling process. By solving the integral of the product of the gamma distribution of abundances and a Poisson sampling distribution, we find that the distribution of read counts of species $i$ in sample $j$ follows a negative binomial [7, 8].

$$P(n_{i,j}|\bar{x}_i, \beta_i, N_j) = \frac{\Gamma(\beta_i + n_{i,j})}{n_{i,j}!\Gamma(\beta_i)} \left(\frac{\bar{x}_i N_j}{\beta_i + \bar{x}_i N_j}\right)^{n_{i,j}} \left(\frac{\beta_i}{\beta_i + \bar{x}_i N_j}\right)^{\beta_i} . \tag{S2}$$

where $N_j$ is the total number of reads at sample $j$. This distribution was used to infer the statistical moments of each ASV that was present in all samples for a given host via maximum likelihood. Specifically, maximum likelihood was numerically performed using `statsmodels v0.14.4` using the empirical distribution of read counts of a given ASV and the total number of reads.

We then leveraged properties of the gamma distribution to evaluate the extent that ASVs with widely varying abundances displayed qualitatively similar sojourn trajectories. Specifically, given that relative abundances follow a gamma distribution with the following shape and rate parameters

$$x_i \sim \text{Gamma}\left(\beta_i, \frac{\beta_i}{\bar{x}_i}\right) \tag{S3}$$

where $\beta_i$ is the squared inverse coefficient of variation

$$\beta_i = \frac{\bar{x}_i^2}{\text{Var}(x_i)} \tag{S4}$$

we can remove the dependency on the mean as follows, reducing the gamma to a single parameter distribution

$$\tilde{x}_i \equiv \frac{x_i}{\bar{x}_i} \sim \text{Gamma}\left(\beta_i, \beta_i\right) \tag{S5}$$

where $\bar{\tilde{x}}_i = 1$. We note that if $\beta_i = 1$ under this rescaling, then $P(\tilde{x}_{i,j}|\beta_i)$ reduces to an exponential distribution. Throughout this study we will be concerned with the natural *logarithm* of relative abundance, requiring us to derive the probability distribution of $\ln \tilde{x}$. Setting $y \equiv \ln \tilde{x}$, we obtain

$$P(y_i|\beta_i, \beta_i) = P(\tilde{x}_i|\beta_i, \beta_i) \cdot \left|\frac{d}{dy_i} e^{y_i}\right| \tag{S6a}$$

$$= \frac{\beta_i^{\beta_i}}{\Gamma\left(\beta_i\right)} \exp\left[\beta_i (y_i - e^{y_i})\right] \tag{S6b}$$

with an expected value

$$\langle y_i \rangle = \psi(\beta_i) - \ln \beta_i \tag{S7}$$

where $\psi(\cdot)$ is the digamma function. Sojourn properties were examined using the rescaled variable $y_i(t) - \bar{y}_i$, where $\bar{y}_i$ is the time-averaged mean which is equal to the ensemble mean $\langle y_i \rangle$ when the dynamics of a community member are stationary and ergodic.

#### Pattern 1: Sojourn time distribution

The relative abundance trajectory of a given ASV $[y_i(t_1), y_i(t_2, ..., y_i(t_M))]$, where $M$ is the number of samples, was converted into a boolean vector representing whether a given observation was greater or less than $\bar{y}_i$ $([+, +, +, ..., -, -])$. The number of steps in a sojourn period was calculated as the number of consecutive observations of the same sign. This number was converted to the number of days by matching sample indices to the sample metadata. Sojourn times of different ASVs belonging to the same host were merged into a single empirical distribution $P(\mathcal{T})$. A null distribution based on the data was obtained by permuting each timeseries and repeating our procedure for calculating sojourn times. This approach accounts for heterogeneity in the time between sampling events.

We note that in principle one could compare empirical distributions *between* different hosts. However, we expect the dynamics of a given ASV to be effectively independent in different hosts, absent a clear mechanism (e.g., immigration between hosts in the same household).

We are also able to derive a prediction for $P(\mathcal{T})$ solely using the observation that $x_i$ tends to follow a gamma distribution, representing the the scenario where the time between sampling events is much larger than the timescale of growth ($\delta t \gg \tau$). In this parameter regime there remains the possibility that both samples will be greater than $\bar{x}_i$ even if two sampling events effectively represent two independent draws from the stationary distribution. The probability of observing $\mathcal{T}$ consecutive draws can be modeled as a geometric process using the cumulative distribution function of the log of our rescaled gamma random variable. Defining $\Phi \equiv \int_{\langle y_i \rangle}^{\infty} P(s|\bar{y}_i, \beta_i) ds$, the probability of obtaining $\mathcal{T}$ consecutive random variables $> \langle y_i \rangle$ is $\Phi^{\mathcal{T}}$. Similarly, the probability of obtaining $\mathcal{T}$ draws $< \langle y_i \rangle$ is $(1-\Phi)^{\mathcal{T}}$. To obtain the full probability distribution it is necessary to identify the normalization constant over the domain $\mathcal{T} \in \{t \in \mathbb{Z} \mid t \geq 1\}$, as we are treating time as a discreet number of days. In this null gamma analysis, we set the upper bound of $\mathcal{T}$ as the number of *samples* collected in a given host. This choice was made because if our samples truly represented draws from a stationary distribution, than the time between samples would be irrelevant.

$$P(\mathcal{T}) = \frac{\Phi^{\mathcal{T}} + (1-\Phi)^{\mathcal{T}}}{Z_{\text{finite}}} \tag{S8a}$$

$$Z_{\text{finite}} = \sum_{t \in \#\,\text{samples}} \left( \Phi^{t} + (1-\Phi)^{t} \right) \tag{S8b}$$

The above null distribution was calculated for each host as a mixture over ASVs, where the weight of each ASV was defined as the fraction of observed sojourn periods belonging to said ASV. Because the above result depends only on the *stationary* distribution it remains valid for sampling events with different time intervals (e.g., most samples taken daily, with some taken ever other day).

#### Pattern 2: Sojourn time vs. sojourn area

We are interested in investigating the relationship between the number of days over which a sojourn trajectory occurs and the area under the sojourn trajectory. For several classes of random walks the sojourn trajectory has been investigated as the mean deviation from the origin for some $t \in [0, \mathcal{T}]$: $\langle x(t) - x(0) \rangle_{\mathcal{T}}$ [9, 10]. The area over sojourn time $\mathcal{T}$ can be defined as an integral, where the relationship between the two variables has the following form

$$\mathcal{A}(\mathcal{T}) \equiv \int_0^1 \langle x(s\mathcal{T}) \rangle_{\mathcal{T}} - x(0) \propto \mathcal{T}^{\alpha} \tag{S9}$$

for unknown exponent $\alpha$. We note that this definition can be interpreted as an area as it represents an integral over a scalar function with one variable ($\mathcal{T}$ is treated as a constant for a given trajectory). In this study we leveraged the observation that ASVs do not tend to go extinct nor take over the community, but that relative abundances fluctuate around an intermediate steady-state abundance. Using the log rescaled relative abundances defined above, we calculated the following integral for all sojourn periods $\mathcal{T} \geq 8$ as

$$\mathcal{A}(\mathcal{T}) \equiv \int_0^1 (y_i(s\mathcal{T}) - \bar{y}_i) ds \tag{S10}$$

We do not consider sojourn trajectories with $< 8$ observations to ensure a reasonable minimum number of timepoints. We note that deviations are never *exactly* equal to zero at the start nor the end of a sojourn period. Rather, it is necessary to choose some threshold value $|\epsilon| \ll 1$ to identify the *beginning* and *end* of each sojourn trajectory, as rare sojourn trajectories that start/end with anomalously large values can bias the resulting integral. Therefore, throughout our empirical analyses and simulations we only consider sojourn walks if they start and end with $(y_i(0) - \bar{y}_i)$ and $(y_i(\mathcal{T}) - \bar{y}_i) < |\epsilon| = 0.01$. Numerical integration was performed with the composite Simpson's rule using `SciPy`.

##### Pattern 3: Sojourn trajectory

Keeping with the theorized scaling relationship, we estimated the mean deviation within a sojourn trajectory for each possible sojourn time [9, 10]. In order to investigate potential differences between sojourn trajectories with different $\mathcal{T}$, we calculated the empirical mean of each $\mathcal{T}$ from sojourn trajectories pooled across ASVs and hosts. We only examined the shape of a given $\mathcal{T}$ if there were at least five timepoints. For completion, individual sojourn trajectories were plotted separately for each host (Fig. S8).

##### Additional temporal patterns

To evaluate the degree that sojourn trajectories reflect temporal dynamics, we examined the difference between empirical distributions of $\mathcal{T}$ and their time-permuted nulls. We then compared that difference to those obtained from two previously characterized macroecological distributions of variables that have the dimension of time. Those two distributions were of the length of time where an ASV was consecutively: 1) observed (i.e., residence time, $t_{\mathrm{res}}$) and 2) unobserved (i.e., return time, $t_{\mathrm{ret}}$). These two quantities were selected because 1) they have the same dimension has $\mathcal{T}$ (i.e., time), 2) their distributions have been an object of study in prior macroecological investigations [11, 12, 13], and 3) the degree that they reflect temporal dynamics has been debated [14, 15].

We define these quantities below. First, we define an indicator function that reflects whether an ASV in sample $m$ is present or absent

$$I_m = \mathbf{1}_{\{x(t_m)>0\}} \tag{S11}$$

The beginning and end of the $k$th trajectory where a given ASV is consistently observed can then be defined as

$$\underline{m}_k, \overline{m}_k = \begin{cases} I_{\underline{m}_k - 1} = 0, \\ I_{\overline{m}_k + 1} = 0, \\ I_m = 1 \quad \text{for all } \ \underline{m}_k \leq m \leq \overline{m}_k \end{cases}$$

from which we define the residence and return times

$$t^k_{\text{res}} \equiv t_{\overline{m}_k} - t_{\underline{m}_k}$$
$$t^k_{\text{ret}} \equiv t_{\underline{m}_{k+1}} - t_{\overline{m}_k}$$

Because $t_{\text{res}}$ and $t_{\text{ret}}$ depend on an ASV being periodically *absent*, these quantities cannot, by definition, be calculated from the set of ASVs used to investigate sojourn trajectories. Therefore, we selected ASVs absent in at least one timepoint with at least 10 non-zero observations. Null distributions were obtained by permuting the empirical read count trajectory of each ASV $10^3$ times. The difference between the empirical and time-permuted null distribution was quantified using Jensen–Shannon divergence.

### Sojourn patterns of the Stochastic Logistic Model

Below we derive predictions and interpretations for the three empirical sojourn patterns using the Stochastic Logistic Model of growth (SLM).

#### General properties

A property of the SLM that makes it a useful ecological model is that its stationary distribution (i.e., distribution of $x(t)$ as $t \to \infty$) is the gamma distribution. Alternatively stated, the SLM predicts the form of the AFD we observe in natural communities. The stationary distribution of the SLM is a gamma distribution, The mean and inverse squared CV of the gamma distribution can be defined in terms of the ecological parameters

$$\bar{x}_i = K_i \left(1 - \frac{\sigma_i}{2}\right) \tag{S12a}$$

$$\beta_i = \frac{2 - \sigma_i}{\sigma_i} \tag{S12b}$$

where we can obtain the SLM for $x_i$ rescaled by $\bar{x}_i$ ($\tilde{x}_i$) as

$$\frac{d\tilde{x}_i}{dt} = \frac{\tilde{x}_i}{\tau_i}\left(1 - \tilde{x}_i\left(1 - \frac{\sigma_i}{2}\right)\right) + \sqrt{\frac{\sigma_i}{\tau_i}}\tilde{x}_i \cdot \eta_i(t) \tag{S13}$$

When one is investigating the natural logarithm of $\tilde{x}_i$ ($y_i$), the equivalent SDE can be derived from SLM by expanding $dy_i$ as a Taylor series and using Itô's lemma.

$$dy_i = \frac{\partial y_i}{\partial t}dt + \frac{\partial y_i}{\partial \tilde{x}_i}d\tilde{x}_i + \frac{1}{2}\frac{\partial^2 y_i}{\partial \tilde{x}_i^2}(d\tilde{x}_i)^2 \tag{S14a}$$

$$= \frac{\partial y_i}{\partial t}dt + \frac{\partial y_i}{\partial \tilde{x}_i}\left(\frac{\tilde{x}_i}{\tau_i}\left(1-\tilde{x}_i\left(1-\frac{\sigma_i}{2}\right)\right) + \sqrt{\frac{\sigma_i}{\tau_i}}\tilde{x}_i \cdot \eta_i(t)\right)$$

$$+ \frac{1}{2}\frac{\partial^2 y_i}{\partial \tilde{x}_i^2}\left(\frac{\tilde{x}_i}{\tau_i}\left(1-\tilde{x}_i\left(1-\frac{\sigma_i}{2}\right)\right) + \sqrt{\frac{\sigma_i}{\tau_i}}\tilde{x}_i \cdot \eta_i(t)\right)^2 \tag{S14b}$$

$$= \left(\frac{\partial y_i}{\partial t} + \frac{\partial y_i}{\partial \tilde{x}_i}\frac{\tilde{x}_i}{\tau_i}\left(1-\tilde{x}_i\left(1-\frac{\sigma_i}{2}\right)\right) + \frac{\sigma_i}{2\tau_i}\frac{\partial^2 y_i}{\partial \tilde{x}_i^2}\tilde{x}_i^2\right)dt$$

$$+ \sqrt{\frac{\sigma_i}{\tau_i}}\frac{\partial y_i}{\partial \tilde{x}_i}\tilde{x}_i dW_i(t) \tag{S14c}$$

where $W_i(t)$ is a Wiener process. We then obtain the Langevin by dividing both sides by $dt$

$$\frac{dy_i}{dt} = \frac{1}{\tau_i}\left(1-\frac{\sigma_i}{2}\right)(1-e^{y_i}) + \sqrt{\frac{\sigma_i}{\tau_i}}\eta_i(t) \tag{S15a}$$

$$\approx -\frac{y_i}{\tau_i}\left(1-\frac{\sigma_i}{2}\right) + \sqrt{\frac{\sigma_i}{\tau_i}}\eta_i(t) \tag{S15b}$$

where we have assumed $y_i \ll 1$. This limit can be interpreted as assuming that the CV of $x_i$ is small, an assumption that is suppurated by the range of CVs inferred from empirical timeseries. The resulting SDE is an Ornstein-Uhlenbeck process [16], the time-dependent solution of which is [17]

$$y_i(t) = y_i(0)e^{-\frac{t}{\tau_i}\left(1-\frac{\sigma_i}{2}\right)} + \sqrt{\frac{\sigma_i}{\tau_i}}\int_0^t e^{-\frac{(t-t')}{\tau_i}\left(1-\frac{\sigma_i}{2}\right)}dW_i(t') \tag{S16}$$

with the corresponding time-dependent probability distribution for initial condition $P(y_i, 0) = \delta(y_i - y_i(0))$

$$P(y_i, t|y_i(0), t_0) = \frac{1}{\sqrt{2\pi\,\mathrm{Var}(y_i(\delta t))}}e^{\frac{-(y_i - \langle y_i(\delta t)\rangle)^2}{2\mathrm{Var}(y_i(\delta t))}} \tag{S17}$$

The mean and variance depend only on the time difference $\delta t \equiv t - t_0$ and are defined as

$$\langle y_i(\delta t)\rangle = y_i(0)\exp\left[-\frac{\delta t(1-\sigma_i/2)}{\tau_i}\right] \tag{S18a}$$

$$\mathrm{Var}(y_i(\delta t)) = \left(\frac{2}{\sigma_i}-1\right)^{-1}\left(1-e^{-\frac{2\delta t(1-\sigma_i/2)}{\tau_i}}\right) \tag{S18b}$$

where the autocovariance between observations separated by time difference $\delta t$ can be found to be

$$\langle y_i(t)y_i(t')\rangle = \left\langle\left(y_i(0)e^{-\frac{t}{\tau_i}\left(1-\frac{\sigma_i}{2}\right)} + \sqrt{\frac{\sigma_i}{\tau}}\int_0^t e^{-\frac{(t-u)}{\tau_i}\left(1-\frac{\sigma_i}{2}\right)}dW_i(u)\right)\right.$$

$$\left.\cdot\left(y_i(0)e^{-\frac{t}{\tau_i}\left(1-\frac{\sigma_i}{2}\right)} + \sqrt{\frac{\sigma_i}{\tau_i}}\int_0^{t'} e^{-\frac{(t'-v)}{\tau_i}\left(1-\frac{\sigma_i}{2}\right)}dW_i(v)\right)\right\rangle$$

The cross-terms cancel out and the final term can be solved using properties of Itô calculus, obtaining

$$\langle y_i(t) y_i(t') \rangle = \left( \frac{2}{\sigma_i} - 1 \right)^{-1} e^{-\frac{|t-t'|}{\tau_i}(1-\frac{\sigma_i}{2})} \tag{S19}$$

from which the autocorrelation can be obtained if the process is stationary

$$\rho(|t-t'|, \tau_i, \sigma_i) \equiv \frac{\langle y_i(t) y_i(t') \rangle}{\sqrt{\mathrm{Var}(y_i(t))\mathrm{Var}(y_i(t'))}} = e^{-\frac{|t-t'|}{\tau_i}(1-\frac{\sigma_i}{2})} \tag{S20}$$

Given the recent application of temporal autocorrelation and cross-correlation measures to microbial community data, it is worth noting that the above autocorrelation result extends to autocorrelations calculated using rescaled log-transformed relative abundances [18]. However, the above result holds for the SLM as a *phenomenological* model in the specified parameter limits, whereas measures like cross-correlation were used to infer the resource usage structure of a community in the context of a *mechanistic* consumer-resource model [18]. The sojourn predictions derived in the subsequent section are calculated for individual community members, meaning that pairwise information is not considered. How our sojourn trajectory work can be extended to consider pairwise information, ideally in the context of a mechanistic model, is an open question of interest. We briefly note that the SLM may not be the sole model that can be reduced to an Ornstein–Uhlenbeck process under certain parameter limits and a log-transformation, meaning that while the SLM has been previously justified as a valid minimal model of microbial ecological dynamics, there potentially exists a wider class of dynamical models that are able to reproduce the observed statistical sojourn properties.

#### Derivation of sojourn predictions

With our time-dependent solution in-hand, we can derive predictions for various aspects of sojourn trajectories. It is important to note that centering the random variable $y_i$ around its expected value allows us to model the sojourn trajectories as a stochastic process that *returns to the origin*. This detail is key for investigating timeseries such microbial communities where taxa tend to persist at intermediate abundances, where the observed presence/absence of a community member is primarily determined by sampling [8]. For convenience, we drop the subscript $i$ from our notation, though we note that all predictions were using ASV-specific inferred estimates of $\sigma_i$ and $K_i$.

#### Pattern 1: Sojourn time distribution

The sojourn time distribution can be obtained from the first-passage time (FPT) distribution. The FPT distribution of the OU process has been derived multiple times through different means [19, 20, 21, 22, 23], most recently through the discovery of the existence of a universality class obtained by space and time transformations [24]. We note that first passage time distributions and various quantities derived from them have been applied to ecological communities in the past (see [25] for a review), though typically they are used to investigate the gain and loss of community members, often due to demographic noise, rather than their stochastic excursions from a non-trivial steady-state abundance.

Due to our initial domain being $x \in (0, \infty)$, the domain of our log-transformed variable $y$ is $(-\infty, \infty)$. Under this new domain we can define the first-passage problem as the time for a random variable $y$ to reach the value of zero given that $y_0 > 0$, a scenario that permits the application of the method of images. Here the probability of observing a sojourn time $\mathcal{T}$ is

$$P(\mathcal{T}|\tilde{\tau},\tilde{y}_0) = \sqrt{\frac{2}{\pi}}\frac{\tilde{y}_0 e^{-\frac{\mathcal{T}}{\tilde{\tau}}}}{(1-e^{-\frac{2\mathcal{T}}{\tilde{\tau}}})^{\frac{3}{2}}} \cdot \exp\left[-\frac{\tilde{y}_0^2 e^{-\frac{2\mathcal{T}}{\tilde{\tau}}}}{2\left(1-e^{-\frac{2\mathcal{T}}{\tilde{\tau}}}\right)}\right] \tag{S21}$$

where $\tilde{y}_0 \equiv y_0\sqrt{\frac{2}{\sigma}-1}$ and $\tilde{\tau} \equiv \tau\left(1-\frac{\sigma}{2}\right)^{-1}$. To translate this first-passage time distribution to a sojourn time distribution it is necessary to investigate its behavior as $y_0 \to 0$, corresponding to $\tilde{y}_0 \to 0$. Thus, the distribution reduces to

$$P(\mathcal{T}|\tilde{\tau},\tilde{y}_0) \approx \sqrt{\frac{2}{\pi}}\frac{\tilde{y}_0 e^{-\frac{\mathcal{T}}{\tilde{\tau}}}}{(1-e^{-\frac{2\mathcal{T}}{\tilde{\tau}}})^{\frac{3}{2}}} \tag{S22}$$

For comparing the predicted distribution to data, it is important to consider that $\tilde{y}_0$ can have an appreciable, though small, value due to our choice of $\epsilon$. Therefore, when comparing $P(\mathcal{T})$ to empirical distributions we used Eq. S21. We also discretized our continuous PDF so that it could be compared with the empirical probability density (technically a probability mass function). Since we observe time in discreet units of days, we integrated Eq. S21 for each value of $\mathcal{T}$.

$$\mathrm{P}(\mathcal{T}|\tilde{\tau},\tilde{y}_0) = \int_{\mathcal{T}}^{\mathcal{T}+1} P(\mathcal{T}'|\tilde{\tau},\tilde{y}_0)d\mathcal{T}' \tag{S23}$$

This distribution was then compared to the empirical distribution. We found that a value of $\tau = 4$ set for all ASVs sufficienctly capatured the bulk of the empirical distribution (Fig. 1b).

In order to assess the variation in sojourn times across community members we examined the predicted mean sojourn time. In this case, $\langle\mathcal{T}\rangle$ can be derived by considering the moment generating function of the first-passage time

$$M_{\mathcal{T}}(t;\tilde{y}_0,\tilde{\tau}) = 1 + \frac{t}{\tilde{\tau}}\frac{2^{\frac{t}{2\tilde{\tau}}}}{\Gamma(1-\frac{t}{2\tilde{\tau}})}\int_0^\infty u^{-\frac{t}{\tilde{\tau}}}e^{-\frac{u^2}{2}}\left[\frac{1-e^{-\tilde{y}_0 u}}{u}\right]du \tag{S24}$$

From which we obtain the first moment

$$\langle\mathcal{T}|\tilde{y}_0,\tilde{\tau}\rangle = \left.\frac{dM_{\mathcal{T}}}{dt}\right|_{t=0} = \tilde{\tau}\int_0^\infty e^{-\frac{u^2}{2}}\left[\frac{1-e^{-\tilde{y}_0 u}}{u}\right]du \tag{S25}$$

which was numerically evaluated for values of $\tau$ and with $\sigma_i$ set by inferred parameters for each ASV. We can also identify the expected value in the limits $\tilde{y}_0 \to \infty, 0^+$, corresponding to the weak and strong noise limiting behavior

$$\langle\mathcal{T}\rangle = \begin{cases}\tilde{\tau}\left[\ln\tilde{y}_0 + \frac{\ln 2+\gamma}{2} + \mathcal{O}(1)\right], & \tilde{y}_0 \to \infty \\ \tilde{\tau}\tilde{y}_0\sqrt{\frac{\pi}{2}}, & \tilde{y}_0 \to 0^+\end{cases} \tag{S26}$$

where $\gamma$ is Euler's constant. Keeping $y_0$ fixed, this parameter limit corresponds to the strength of environmental noise approaching zero. Given that the upper bound of $y_0$ is set by our chosen value of $\epsilon = 0.1$ and the median $\sigma_i$ across ASVs is $\approx 0.54$, we expect $\tilde{y}_0 \sim \mathcal{O}(10^{-1})$ which would place the parameter regime closer to the strong noise limit. We note that $\langle\mathcal{T}\rangle$ is not a function of the carrying capacity $K_i$, meaning that we cannot define it as a function of $\bar{x}_i$. This result implied the *absence* of a relationship between $\langle\mathcal{T}\rangle$ and $\bar{x}_i$. However, because the CV is solely a function of $\sigma_i$ we can define $\langle\mathcal{T}\rangle$ as a function of the CV, predicting the existence of a relationship between the two quantities. This is a result that holds at the level of individual ASVs, meaning that its validity holds regardless of the relationship between between statistical moments of abundance *across* ASVs (e.g., Taylor's Law [26]). We note that under a strict interpretation of Taylor's Law the CV will be independent of the mean relative abundance, a prediction that holds in all but one host (Fig. S9).

To compare predictions to data we used the discretized form

$$\langle \mathcal{T} | \tilde{y}_0, \tilde{\tau} \rangle = \sum_{t=1}^{T_{\max}} t \cdot \mathrm{P}(t|\tilde{\tau}, \tilde{y}_0) \tag{S27}$$

where $T_{\max}$ was the total number of days for a given time series. This discretized distribution was normalized, as sojourn periods greater than $T_{\max}$ cannot be observed by definition.

#### Pattern 2: Sojourn time vs. sojourn area

To calculate the expected position of a community member during a sojourn period, it is necessary to first obtain the probability that a walk originating at $y_0$ at time 0 reaches $y$ at time $t$. The Ornstein–Uhlenbeck process is linear, allowing for the application of the image method to Eq. S17 [27].

$$P(y, t|y_0, 0) = \frac{1}{\sqrt{2\pi \,\mathrm{Var}(y(t))}} \left[ e^{\frac{-(y - \langle y(t) \rangle)^2}{2\mathrm{Var}(y(t))}} - e^{\frac{-(y + \langle y(t) \rangle)^2}{2\mathrm{Var}(y(t))}} \right] \tag{S28}$$

as $y_0 \to 0$, the distribution reduces to

$$P(y, t|y_0, 0) = \frac{2y \,\langle y(t) \rangle}{\sqrt{2\pi \,\mathrm{Var}(y(t))^3}} e^{-\frac{y^2}{2\mathrm{Var}(y(t))}} \tag{S29a}$$

$$= \frac{2yy_0 \exp\left[-\frac{t}{\tilde{\tau}}\right]}{\sqrt{2\pi \,\mathrm{Var}(y(t))^3}} e^{-\frac{y^2}{2\mathrm{Var}(y(t))}} \tag{S29b}$$

By repeating the above calculation we can obtain the probability of *returning* to the origin after a sojourn time of $\mathcal{T}$

$$P(y_0, \mathcal{T}|y, t) = \frac{2yy_0 \exp\left[-\frac{(\mathcal{T}-t)}{\tilde{\tau}}\right]}{\sqrt{2\pi \,\tilde{V}^3(\mathcal{T} - t)}} \cdot \exp\left[ -\left( \frac{y^2}{2} e^{-\frac{2(\mathcal{T}-t)}{\tilde{\tau}}} \cdot \mathrm{Var}(y(\mathcal{T} - t)) \right) \right] \tag{S30}$$

where we have defined a form of the variance that preserves time-reversal symmetry for the parameter $\tilde{\tau}$.

$$\tilde{V}(t) \equiv \mathrm{Var}(t|\tilde{\tau}) e^{\frac{2t}{\tilde{\tau}}} = \mathrm{Var}(t| - \tilde{\tau}) \tag{S31}$$

We can now derive the distribution of the position of a community member at time $t$ within a sojourn period of time $T$

$$\Omega(y, t|y_0, 0; y_0, \mathcal{T}) \equiv P(y, t|y_0, 0) \cdot P(y_0, \mathcal{T}|y, t) \tag{S32a}$$

$$\propto e^{-\frac{T}{\tilde{\tau}}} \left[ \mathrm{Var}(t|\tilde{\tau}) \,\mathrm{Var}(\mathcal{T} - t| - \tilde{\tau}) \right]^{-\frac{3}{2}} \cdot \exp\left[ -\frac{y^2}{2V_{\mathrm{eq}}(t, \mathcal{T})} \right] \tag{S32b}$$

where

$$V_{\mathrm{eq}}(t,\mathcal{T}) \equiv \left[(\mathrm{Var}(t|\tilde{\tau}))^{-1} + (\mathrm{Var}(\mathcal{T}-t|-\tilde{\tau}))^{-1}\right]^{-1} \tag{S33a}$$

$$= \left(\frac{2}{\sigma}-1\right)^{-1}\left[\frac{(1-e^{-\frac{2t}{\tilde{\tau}}})(1-e^{-\frac{2(\mathcal{T}-t)}{\tilde{\tau}}})}{1-e^{-\frac{2\mathcal{T}}{\tilde{\tau}}}}\right] \tag{S33b}$$

Using the definition of average position during a sojourn period [9, 10]

$$\langle y(t)\rangle_{\mathcal{T}} \equiv \lim_{y_0\to 0^+}\frac{\int_0^\infty y\Omega(y,t|y_0,0,y_0,\mathcal{T})dy}{\int_0^\infty \Omega(y,t|y_0,0,y_0,\mathcal{T})dy} \tag{S34}$$

we see that the calculation reduces to an integral over an exponential of Gaussian form, the solution of which is

$$\langle y(t)\rangle_{\mathcal{T}} = \sqrt{\frac{8}{\pi}}\sqrt{\left(\frac{2}{\sigma}-1\right)^{-1}\left[\frac{(1-e^{-\frac{2t}{\tilde{\tau}}})(1-e^{-\frac{2(\mathcal{T}-t)}{\tilde{\tau}}})}{1-e^{-\frac{2\mathcal{T}}{\tilde{\tau}}}}\right]} \tag{S35}$$

Because the solution has a characteristic timescale the shape of the expected sojourn trajectory depends on $\tilde{\tau}$. The equation reduces in two asymptotic limits

$$\langle y(t)\rangle_{\mathcal{T}} = \begin{cases} \sqrt{\frac{8}{\pi}}\sqrt{\frac{t(\mathcal{T}-t)}{\mathcal{T}}}, & t, \mathcal{T}-t \ll \tilde{\tau} \\ \sqrt{\frac{8}{\pi}}\left(\frac{2}{\sigma}-1\right)^{-\frac{1}{2}}, & t, \mathcal{T}-t \gg \tilde{\tau} \end{cases} \tag{S36}$$

The expected area under the sojourn trajectory can be obtained by taking the definite integral of the above result over relative time, from which we obtain

$$\langle \mathcal{A}(\mathcal{T})\rangle = \int_0^1 \langle y(s\cdot\mathcal{T})\rangle_{\mathcal{T}}\, ds = \begin{cases} \sqrt{\frac{\pi}{8}}\mathcal{T}^{\frac{1}{2}}, & t, \mathcal{T}-t \ll \tilde{\tau} \\ \sqrt{\frac{8}{\pi}}\left(\frac{2}{\sigma}-1\right)^{-\frac{1}{2}} = \sqrt{\frac{8}{\pi}}\cdot \mathrm{CV}, & t, \mathcal{T}-t \gg \tilde{\tau} \end{cases} \tag{S37}$$

This result means that when the time between observations within a sojourn period exceeds the timescale of growth we predict that the area will be independent of $\mathcal{T}$. We note that neither of these two parameter regimes depend on the value of $\tau$.

#### Pattern 3: Sojourn trajectory

In the previous sub-sections we demonstrated how the statistical excursions of microbial community members from their typical abundance can be investigated using the tools from random walk theory by first identifying a reasonable SDE of ecological dynamics and then performing an appropriate rescaling of the data. Here we briefly discuss how sojourn trajectories observed in microbial communities relate to the forms. It was previously reported that the sojourn trajectories of a wide range of random walks, including those that are biased, exhibit Levy flights, and have either long or short-range correlations, all exhibit the same scaling [9]

$$\langle y(t)-\bar{y}\rangle_{\mathcal{T}} \propto \mathcal{T}^{\alpha} f(t/\mathcal{T}) \tag{S38}$$

This scaling will hold if $P(\mathcal{T})$ decays algebraically and the exponent $\alpha$ can be interpreted as the time exponent that controls the expected squared deviation from the origin (i.e., the full trajectory of abundances over time) $\left\langle [y(t)-y(0)]^2\right\rangle \simeq t^{\alpha}$. If this scaling relation holds, then rescaled sojourn trajectories will collapse on a single curve once $\alpha$ is identified. We found that $\alpha \approx 0$ for microbial communities in the human gut, corresponding to the $t, \mathcal{T} - t \gg \tilde{\tau}$ limit of Eq. S37. This result implies that no rescaling is required, reducing the relationship to

$$\langle y(t) - \bar{y} \rangle_{\mathcal{T}} \propto f(t/\mathcal{T}) \tag{S39}$$

In this result $\mathcal{T}$ only contributes in a dimensionless form, meaning that there is no explicit dependence on the value of $\mathcal{T}$.

24. Costantino Di Bello, Édgar Roldán, and Ralf Metzler. Universal class of exactly solvable diffusions from space-time transformations, March 2025.

25. Sandro Azaele, Samir Suweis, Jacopo Grilli, Igor Volkov, Jayanth R. Banavar, and Amos Maritan. Statistical mechanics of ecological systems: Neutral theory and beyond. *Reviews of Modern Physics*, 88(3):035003, July 2016.

26. L. R. Taylor. Aggregation, Variance and the Mean. *Nature*, 189(4766):732–735, March 1961. Publisher: Nature Publishing Group.

27. Sidney Redner. *A Guide to First-Passage Processes*. Cambridge University Press, 1 edition, August 2001.

371 # Figures

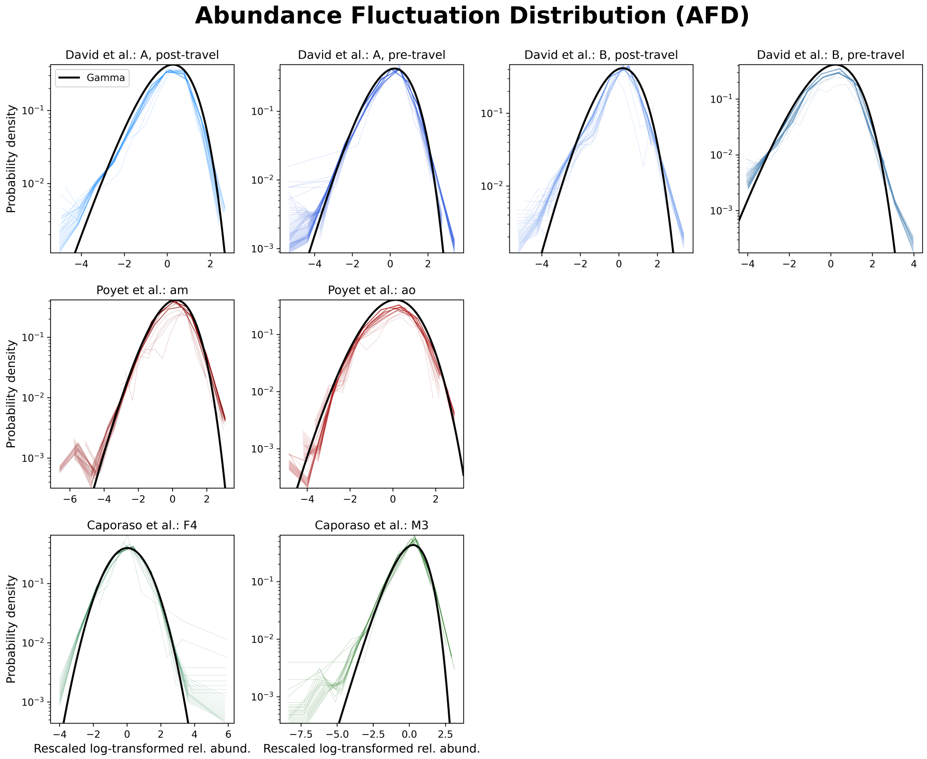


**Figure S1. Empirical temporal AFDs are gamma distributed.** Empirical Abundance Fluctuation Distributions tend to follow a gamma distribution across hosts as well as datasets.

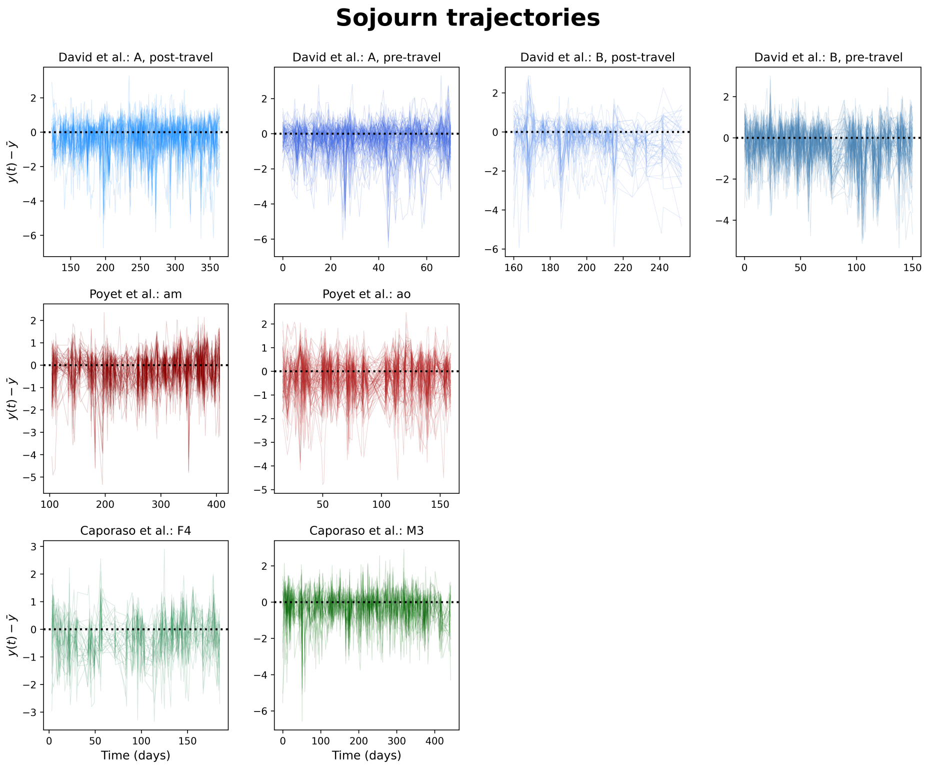


**Figure S2. Deviations of the $\log_e$ rescaled abundance around the time-averaged mean.** A visualization of the trajectories of the deviations of the $\log_e$ rescaled relative abundance around its time-averaged mean. Each line represents one ASV.

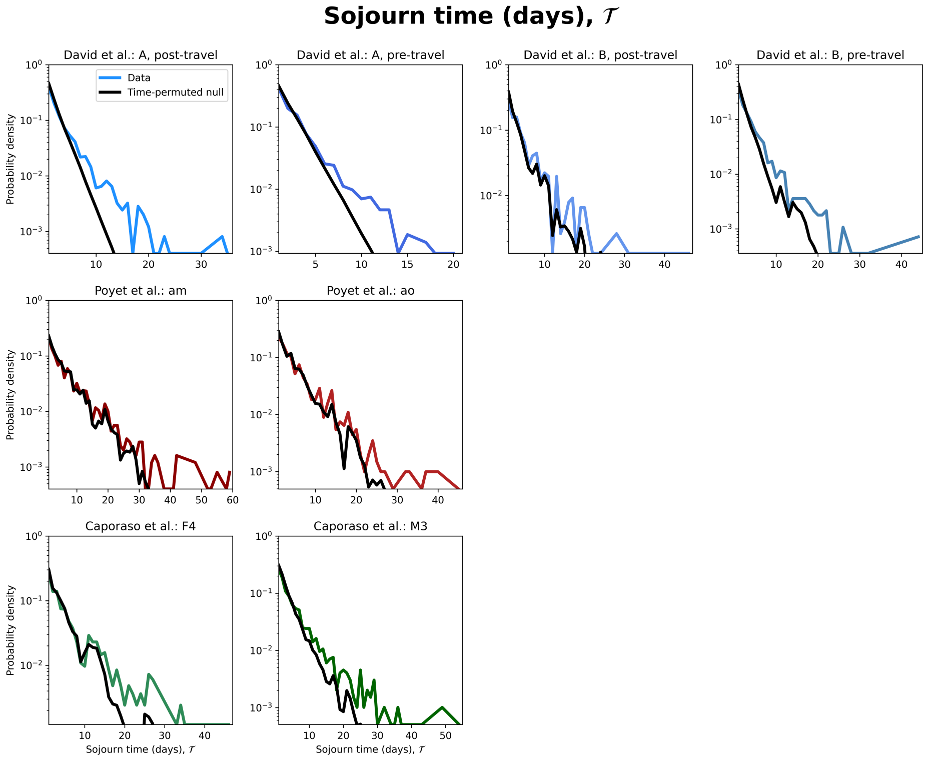


**Figure S3. Empirical sojourn time distributions are not captured by a permutation-based null.** Empirical sojourn distributions for each timeseries display consistent deviations from a null distribution calculated by permuting ASV abundances with respect to time. Null distributions were calculated from $10^3$ permutations.

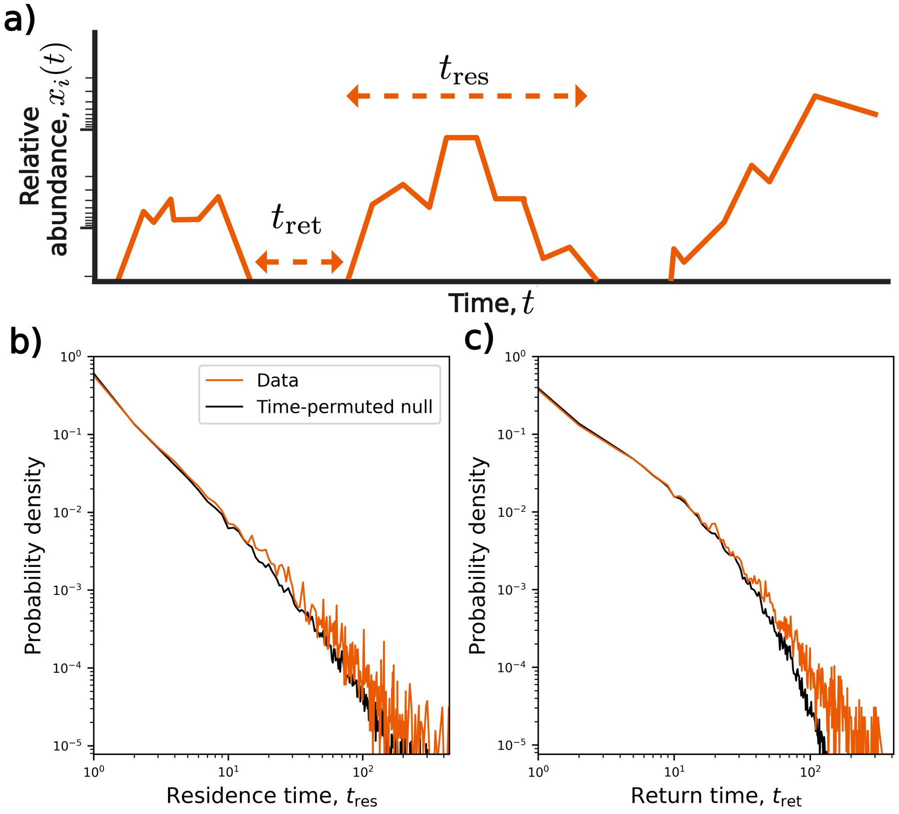


**Figure S4. Residence and return time distributions resemble time-permuted null distributions. a)** We calculated the number of consecutive days where an ASV was observed (residence time, $t_{\mathrm{res}}$) or unobserved (return time, $t_{\mathrm{ret}}$) in each host. **b,c)** An across-hosts mixture distribution was calculated for each measure and a corresponding time-permuted null distribution was obtained. In general the null distributions resembled the empirical distributions. Null distributions were calculated from $10^3$ permutations.

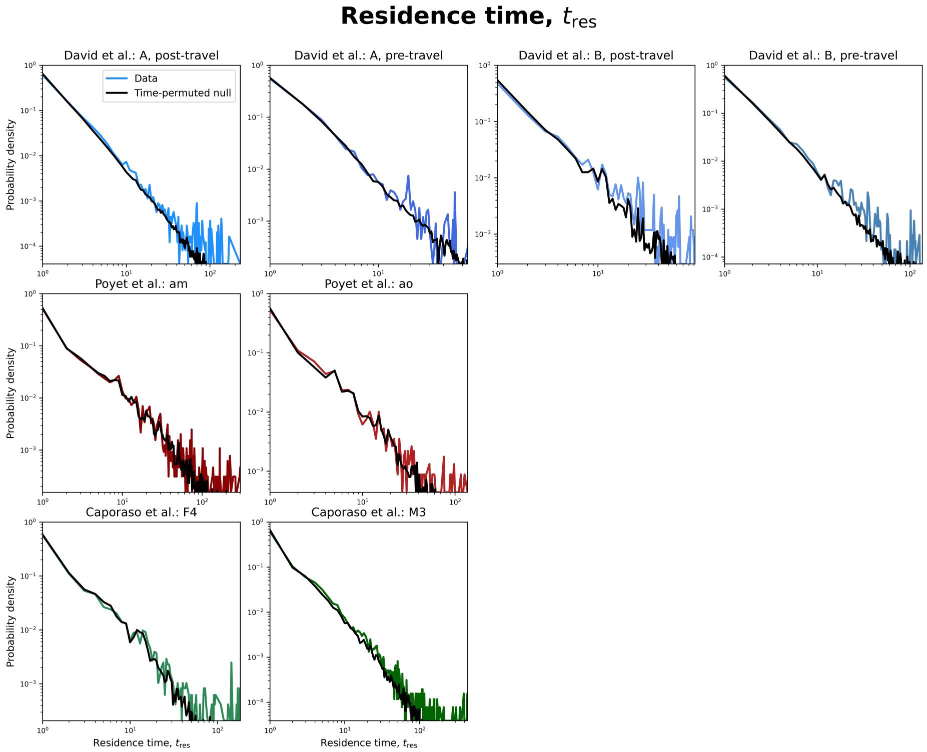


**Figure S5. Empirical and null residence time distributions for separate hosts.** The residence time data presented in Fig. S4 separated by host.

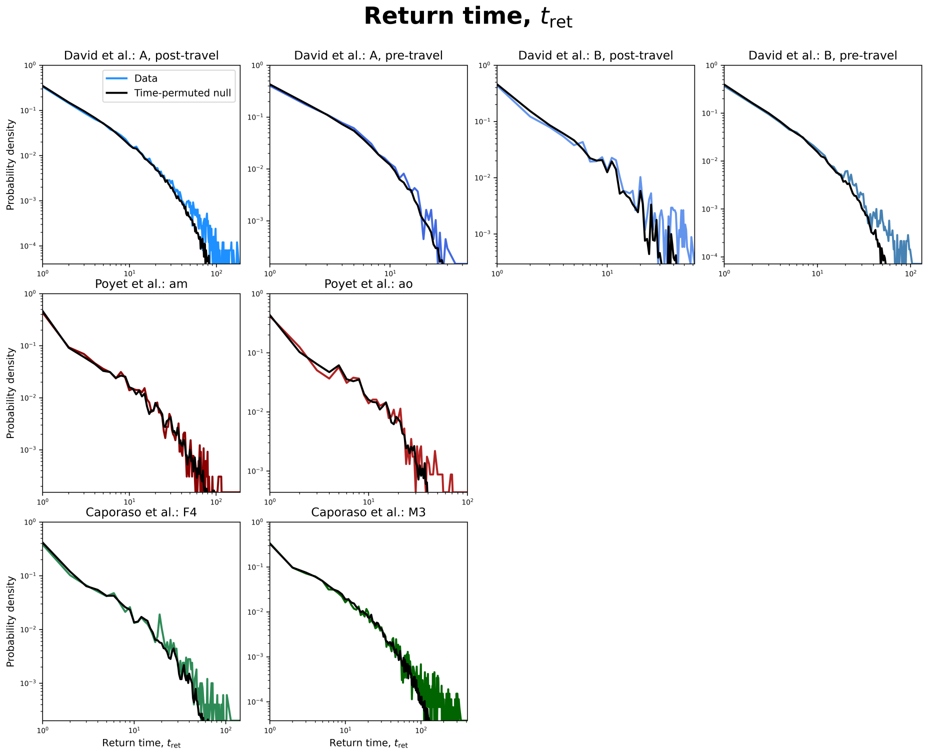


**Figure S6. Empirical and null return time distributions for separate hosts.** The return time data presented in Fig. S4 separated by host.

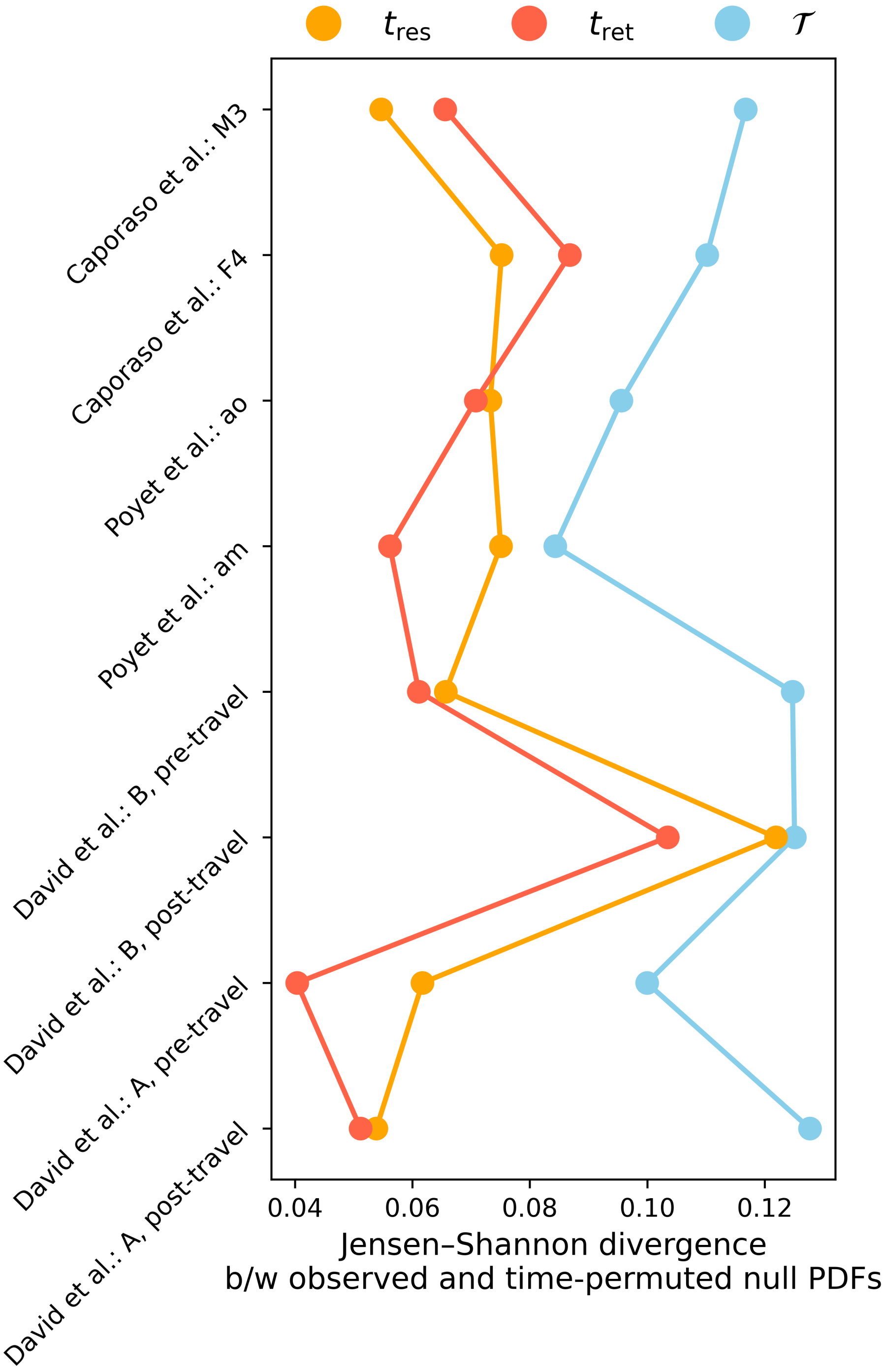


**Figure S7. Sojourn time distributions contain greater temporal information.** Empirical sojourn ($\mathcal{T}$), residence ($t_{\rm res}$), and return time ($t_{\rm ret}$) distributions were compared to their corresponding time-permuted null distributions on a per-host basis. The Jensen-Shannon divergence was used to assess the degree that empirical distributions deviated from the null, where sojourn time distributions consistently displayed greater divergence across all hosts.

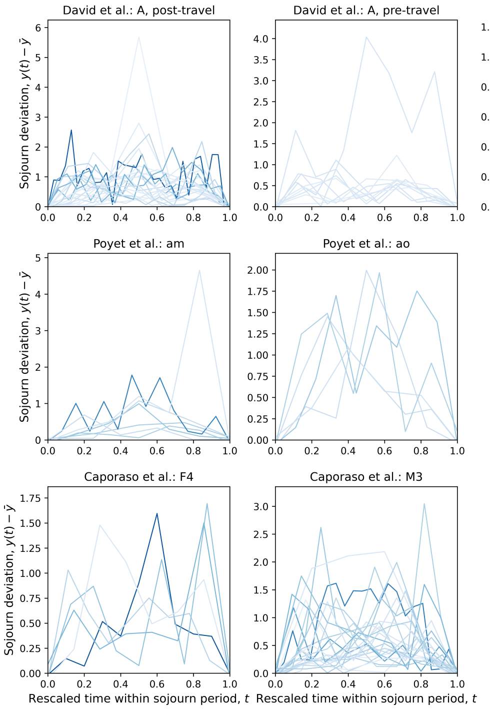


**Figure S8. Individual sojourn trajectories of all hosts and datasets.** Each sojourn trajectory used in Fig. 1d is plotted for a specific host × dataset combination.

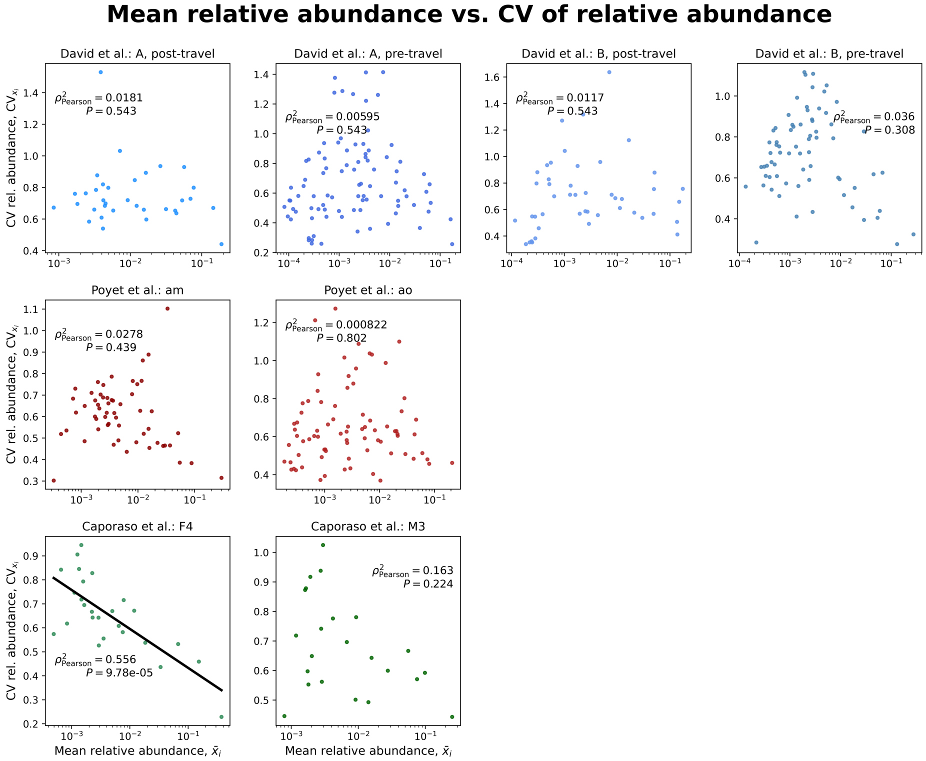


**Figure S9. The CV of relative abundance is generally independent of the mean.** The relationship between the CV of relative abundance and the $\log_{10}$-transformed mean relative abundance was assessed using least-squares regression. There was the absence of a significant relationship in all but a single host. The false discovery rate of the separate regressions was accounted for using the Benjamini–Hochberg procedure.

### Tables

| Reference | Host | Trajectory | # samples | # days | Mean # days b/w samples | # ASVs examined |
|---|---|---|---|---|---|---|
| [1] | F4 | Entire | 88 | 182 | 2.09 | 26 |
| | M3 | Entire | 251 | 442 | 1.77 | 23 |
| [2] | am | Entire | 112 | 302 | 2.72 | 56 |
| | ao | Entire | 65 | 143 | 2.23 | 79 |
| [3] | A | Post-travel | 216 | 241 | 1.12 | 34 |
| | | Pre-travel | 64 | 70 | 1.11 | 90 |
| | B | Post-travel | 54 | 92 | 1.74 | 46 |
| | | Pre-travel | 119 | 150 | 1.27 | 70 |

**Table S1.** Metadata of the timeseries used in this study from reprocessed public datasets. ”Trajectory” refers to whether the entire timeseries was split into multiple, smaller timeseries that met our criteria. The mean number of days between sampling events over all rows weighed by number of samples and number of ASVs is $\approx 1.92$.
